# Thermodynamics of Indirect Readout in Cre-*loxP* Recognition

**DOI:** 10.64898/2026.05.05.721372

**Authors:** Jonathan S. Montgomery, Megan E. Judson, Robert M. Bauer, Matthew J. Benedek, Caroline E. Hervey, Mark P. Foster

## Abstract

Cre, a site-specific tyrosine DNA recombinase, enables targeted manipulation of genetic material without production of cytotoxic double-strand DNA breaks. Though widely deployed in biotechnology, an incomplete understanding of how Cre recognizes its cognate *loxP* target limits its broader application in human health. Cre has been proposed to recognize an inverted pair of recombinase binding elements (RBEs) at *loxP* sites using an indirect readout mechanism, inducing conformations that are disfavored at noncognate sites. Despite a high degree of specificity, structural studies have shown that Cre protomers make few direct base-specific contacts to each RBE, implicating noncontacted positions in recognition by tuning flexibility and enabling binding-coupled conformational changes. We designed a set of *loxP* half-site variants predicted to rigidify the DNA substrate and measured the thermodynamics of Cre binding by isothermal titration calorimetry. Thermodynamic signatures and NMR spectra reveal that unfavorable mutations at noncontacted positions in the RBE reduce binding- coupled conformational changes. Moreover, mass photometry of Cre binding to oligonucleotides containing two properly spaced RBEs revealed that non-contacted positions in the intervening 8-base pair spacer influence cooperative dimerization of two Cre protomers at *loxP* sites, and their synapsis to form catalytically active Cre_4_-*loxP*_2_ complexes. These results demonstrate that noncontacted positions contribute to specificity by encoding favorable DNA mechanics, offering new design principles for engineering Cre variants that target alternative DNA sequences.

## Introduction

The ability to manipulate genetic material presents the opportunity to treat genetic diseases by excision and reintroduction of specific DNA sequences. Nuclease-based gene editing technologies such as CRISPR/Cas9 produce double-strand DNA (dsDNA) breaks at a site of interest that are repaired via mutagenic host repair pathways such as nonhomologous end joining.^1,2^ Tyrosine site-specific DNA recombinases (YSSR), on the other hand, recombine DNA at paired homologous DNA sequences, catalyzing excision, inversion or exchange of genetic material without producing dsDNA breaks.^3,4^ This capacity to preserve genome integrity makes YSSRs attractive tools for therapeutic gene editing.

Cre (<u>C</u>auses <u>Re</u>combination) is the most widely studied YSSR.^3,5^ It has demonstrated utility in laboratory settings to produce conditional knockouts in transgenic mice^6^ and to introduce genes through recombinase mediated-cassette exchange.^7^ Cre is monomeric in solution and comprises two independently folding domains, an N-terminal Core Binding domain (CreCB) and a C-terminal Catalytic domain (CreCat), that encircle the DNA forming a C-shaped clamp (Figure 1 b-c).^8,9^ Cre recombines *loxP* sequences that contain two recombinase binding elements (RBEs) oriented as inverted repeats flanking an 8-base pair asymmetric spacer sequence (Figure 1 a). Binding of two Cre monomers to the RBEs of a *loxP* site proceeds cooperatively, followed by synapsis of two Cre_2_-*loxP* dimers to form the tetrameric Cre_4_-*loxP*_2_ synaptic complex that is active for recombination.^9,10^

**Figure 1.**
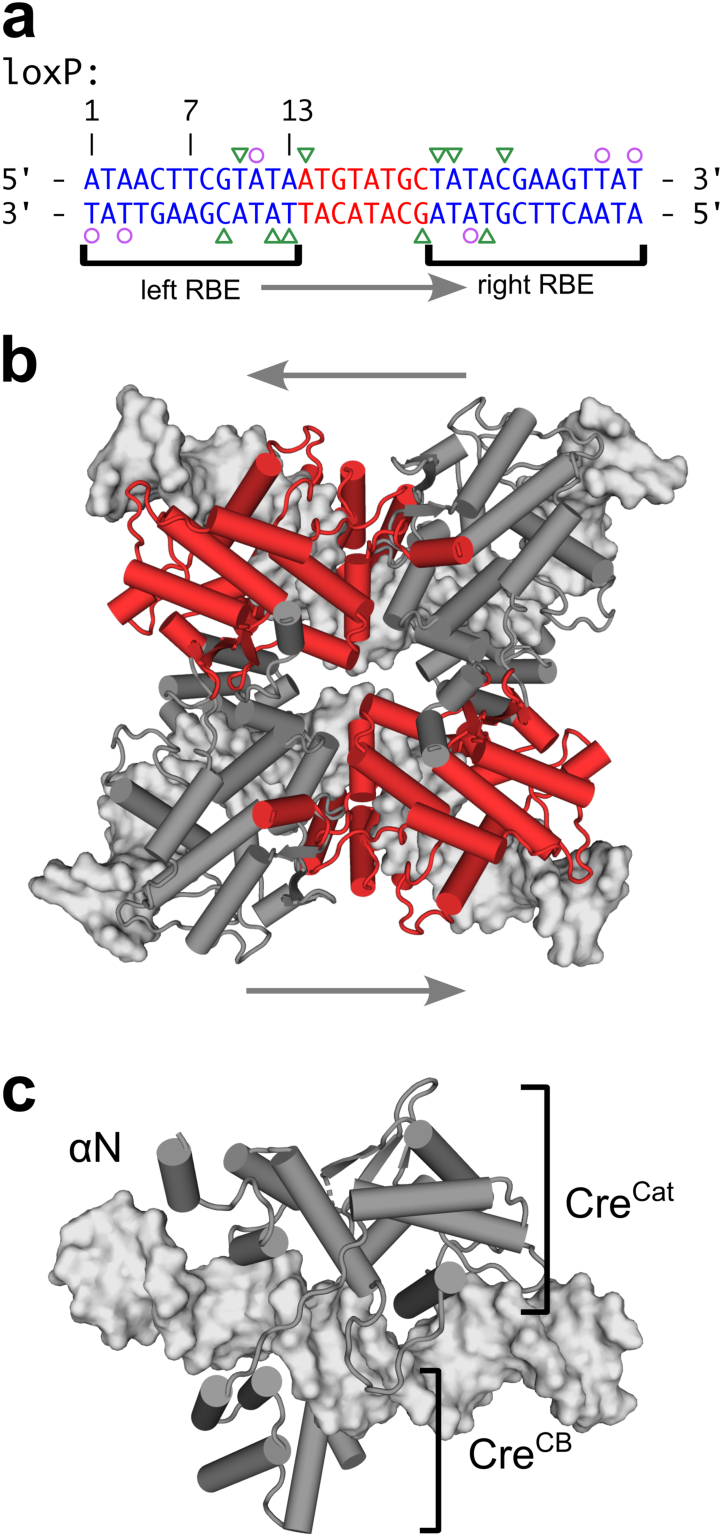
Assembly of Cre-*loxP* complexes is coincident with deformations in *loxP* DNA. (a) *loxP* DNA sequence comprising two inverted recombinase binding elements (RBEs, blue) flanking an asymmetric spacer (red). Triangles and circles denote direct major and minor groove contacts to DNA bases observed in high-resolution structures. Arrows indicate direction of the asymmetric spacer sequence. (b) Tetrameric Cre4-*loxP*2 synaptic complex (PDB 2HOI) with alternating protomers active for single strand DNA cleavage (red). (c) Cre bound as a monomer to a *loxP* half site (PDB 7RHY) features a ∼18° bend in the DNA.

While Cre has demonstrated utility in manipulating genomes, overcoming its obligate pairing with *loxP* sites would broaden its utility for gene editing. Despite high-resolution structures of from x-ray crystallography of Cre bound in tetrameric structures with DNA,^3,11^ and multiple studies that evaluate effect of DNA sequence on recombination,^12–14^ these have been of limited utility in guiding mutagenesis efforts to engineer Cre variants that effectively recombine alternative sites. The most successful approaches for altering Cre’s specificity have implemented substrate linked directed evolution.^15–17^ However, these approaches are laborious and highly empirical and have yet to produce a general framework for understanding how DNA sequence recognition is achieved by Cre and its variants.^3^ Notably, because recombination is an ordered multistep process, protein and DNA mutations may affect multiple steps in the recombination pathway, including (1) the monomer-site recognition, (2) assembly of higher order oligomeric states, (3) DNA cleavage kinetics, and (4) direction of resolution of recombination intermediates.

With increased interest in retargeting Cre to biomedically relevant DNA sequences,^18,19^ understanding the basis of DNA site recognition by Cre gains increased urgency. Broadly, the structural basis of sequence-specific DNA binding by proteins is described as arising from a combination of *direct* and *indirect* readout. Direct readout features non-covalent interactions between specific amino acid side chains and nucleotide-specific chemical groups on the DNA bases, whereas indirect readout is achieved through sequence-specific differences in DNA shape.^20^ Studies focused on recognition of a *loxP* half-site (avoiding dimerization) have shown that site recognition is coupled to deformations in the cognate DNA substrate that are not induced when binding non-cognate sites.^9,21^ Given that Cre makes only three direct base-specific contacts to *loxP* half-sites (Figure 1)^9,10^, yet binds with thousand-fold specificity over noncognate DNA,^21^ this indicates that noncontacted positions play an essential role in defining Cre-*loxP* recognition.

In this report, we test the hypothesis that deleterious mutations at noncontacted positions in the *loxP* sequence function by disrupting indirect readout by Cre. We employed isothermal titration calorimetry (ITC) and nuclear magnetic resonance (NMR) spectroscopy to characterize how noncontacted positions in the *loxP* sequence influence binding thermodynamics and binding-coupled spectral signatures. We observe that mutations in the RBE that disfavor groove distortions exhibit less favorable binding but with paradoxically more favorable binding enthalpy. NMR spectra reveal that mutations at noncontacted RBE positions abrogate protein spectral signatures associated with cognate DNA binding. Lastly, we found that mutations at noncontacted positions in the *loxP* spacer don’t affect monomer-RBE binding but greatly perturb subsequent steps in assembly of Cre-*loxP* synaptic complexes. These results demonstrate that noncontacted positions contribute to specificity by encoding favorable DNA mechanics, offering new design principles for engineering Cre variants that target alternative DNA sequences.

## Results

### Design of loxP half site variants

To probe the role of noncontacted positions in recognition of *loxP* substrates, we generated a set of *loxP* half site variants (comprising an RBE and half of the asymmetric spacer: *loxP*21L, *loxP*21R, *loxP*21L-A11T, *loxP*21L-G16A and loxScramble, Figure 2) intended to alter DNA flexibility and bending. The *loxP*21L and *loxP*21R oligos test whether the spacer sequence affects RBE binding by Cre monomers. To test the importance of groove distortions observed in Cre-bound *loxP* structures, we designed two *loxP* half site variants, A11T and G16A, that introduce a rigid dA-dT base step at positions observed to undergo minor groove distortion upon protein binding (Figure S1 and Table S1).^20,22–25^ The loxScramble mutant was designed to replace all noncontacted positions and exhibit predicted minor groove widths that resemble B-form DNA.^22^

**Figure 2.**
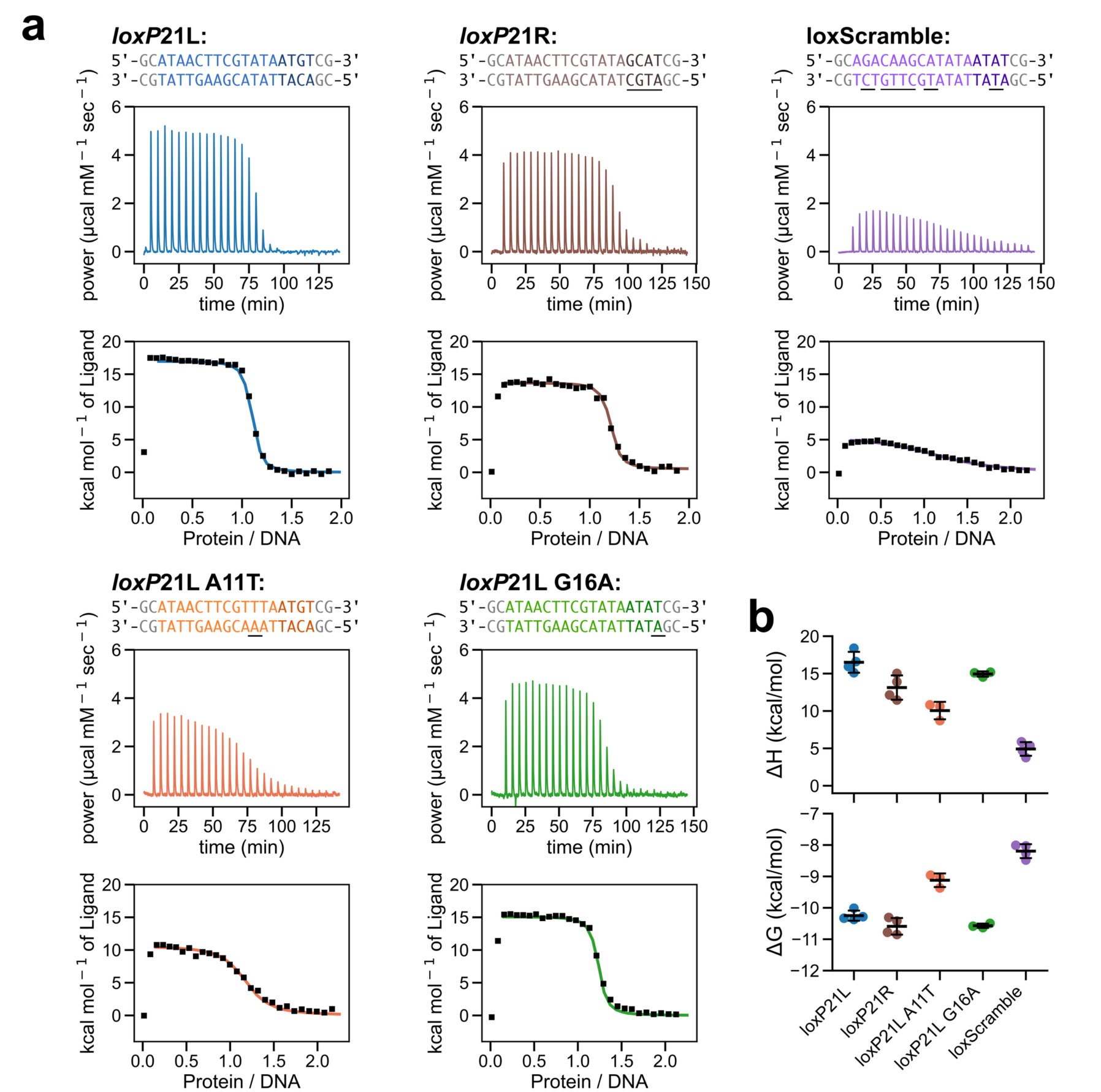
ITC of Cre binding to *loxP* half site variants. (a) Baseline corrected thermograms for titrations with each DNA substrate (top) and integrated heats fit to a single site binding model (bottom). Positions of mutations in *loxP* half-site variants are underlined. (b) Enthalpy (ΔH, top) and free energy of binding (ΔG, bottom) values obtained from fitting titrations to a single site binding model. Plots indicate the mean and standard deviation of replicates.

### Thermodynamics of recognition of loxP half site mutants

We performed isothermal titration calorimetry (ITC) experiments to quantify the effect of noncontact positions on the thermodynamics of DNA half-site binding by Cre. Solutions containing Cre were titrated into solutions containing one of the *loxP* half site variants. As observed with the native *loxP* half site^21^, all titrations exhibited large positive (unfavorable) enthalpies (ΔH) of binding and thermograms were well described by a one site binding model, with small variations from 1:1 stoichiometry attributed to uncertainties in component concentrations (Figure 2). Integrated thermograms were fit using a single site binding model to obtain the binding free energy, enthalpy and entropy.

We tested whether the spacer sequence affects thermodynamics of RBE recognition by Cre. Crystallographic and cryo-EM structures indicate that the first position of the spacer is involved in a nonspecific contact by Cre, and spacers symmetrized to match the left side of the asymmetric half-site have been reported to exhibit increased recombination efficiency relative to right symmetric spacers^26^. To assess whether the spacer affects RBE binding, we performed ITC experiments of Cre binding to half-sites containing either the left or right spacer (*loxP*21L, *loxP*21R; Figure 2 a). Fitting titrations for *loxP*21R yielded a K_D_ of 9.1 ± 4 nM, ΔH of 13.73 ± 0.72 kcal mol^-1^, and TΔS of 24 ± 0.72 kcal mol^-1^ at 15 °C (Figure 2 b). This result is indistinguishable from that of binding to *loxP*21L (p-values of 0.07 and 0.01 for ΔG and ΔH respectively), indicating that the spacer sequence does not contribute to the large unfavorable enthalpy of binding that coincides with high-affinity recognition of *loxP* half site sequences.

Mutations to non-contact positions intended to disfavor DNA bending strongly disfavor recognition be Cre. Titrations with the A11T variant resulted in an elevated K_D_ of 120 ± 40 nM, a less unfavorable ΔH of 10 ± 1 kcal mol^-1^, and favorable TΔS of 19 ± 1 kcal mol^-1^ at 15 °C (p-value < 0.01 for both values, Figure 2 b). Fitting titrations for the G16A variant yielded a K_D_ of 9 ± 1 nM, ΔH of 14.9 ± 0.4 kcal mol^-1^, and TΔS of 25.5 ± 0.4 kcal mol^-1^ at 15 °C. This results in no significant difference in binding affinity or enthalpy compared to the native *loxP* half sites (p-values of 0.02 and 0.12 for ΔG and ΔH respectively). Titrations with the loxScramble variant, which features at non-contact positions nucleotides that favor B-form helical structure, resulted in a much weaker K_D_ of 660 ± 200 nM, ΔH of 4.7 ± 0.8 kcal mol^-1^, and TΔS of 12.9 ± 0.8 kcal mol^-1^ at 15 °C (p-value < 0.01 for both variables). The results underscore the importance of non-contact positions in defining the overall thermodynamics of Cre-DNA recognition.

We performed additional experiments to test whether the experimentally observed large unfavorable enthalpy of Cre-DNA binding is confounded by buffer effects. We performed additional titrations of Cre into loxP21L in buffers with a range of ionization enthalpies (HEPES; 5.0 kcal mol^-1^, ACES; 7.3 kcal mol^-1^, and Tris; 11.4 kcal mol^-1^)^27^ (Figure S2). By considering that observed enthalpy changes include a contribution from transfer of protons to or from the buffer, buffer-independent enthalpies obtained from a linear fit to: 𝛥𝐻_𝑜𝑏𝑠_ = 𝛥𝐻_𝑏𝑖𝑛𝑑_ + 𝑛_𝐻(_ · 𝛥𝐻_𝑖𝑜𝑛_, where ΔH_obs_ is the measured enthalpy of binding, ΔH_bind_ is buffer corrected enthalpy of binding, n_H+_ is the average number of protons transferred from buffer to the macromolecules, and ΔH_ion_ is the enthalpy of ionization for the buffer. This yielded a buffer corrected enthalpy of binding of +14.9 kcal/mol at 15 °C, with 0.6 ± 0.3 ionization events at pH 7.0. This requires only a modest correction to the large unfavorable enthalpy of binding, consistent with large binding-coupled protein-DNA conformational changes.^28^

### Thermodynamic signature of cognate Cre-loxP recognition from temperature-dependent binding enthalpy

Large negative changes in heat capacity, Δ*C_p_*, in excess of that arising from changes in solvation from burial of intermolecular surface area, provides a thermodynamic signature of coupled conformational changes during specific protein-DNA recognition.^29,30^ We quantified this thermodynamic parameter from the linear relation 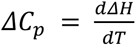 by recording ITC experiments over a range of temperatures from 283 K to 298 K (Figure 3). Binding between Cre and the native *loxP*21L yielded the largest negative Δ*C_p_* (-830 ± 90 cal mol^-1^ K^-1^), whereas for each variant a much smaller value was obtained (*loxP*21L A11T, Δ*C_p_* = -260 ± 70 cal mol^-1^ K^-1^; loxScramble, Δ*C_p_* = - 120 ± 60 cal mol^-1^ K^-1^) (Figure 3b). Binding free energies were relatively insensitive to temperature over this range.

**Figure 3.**
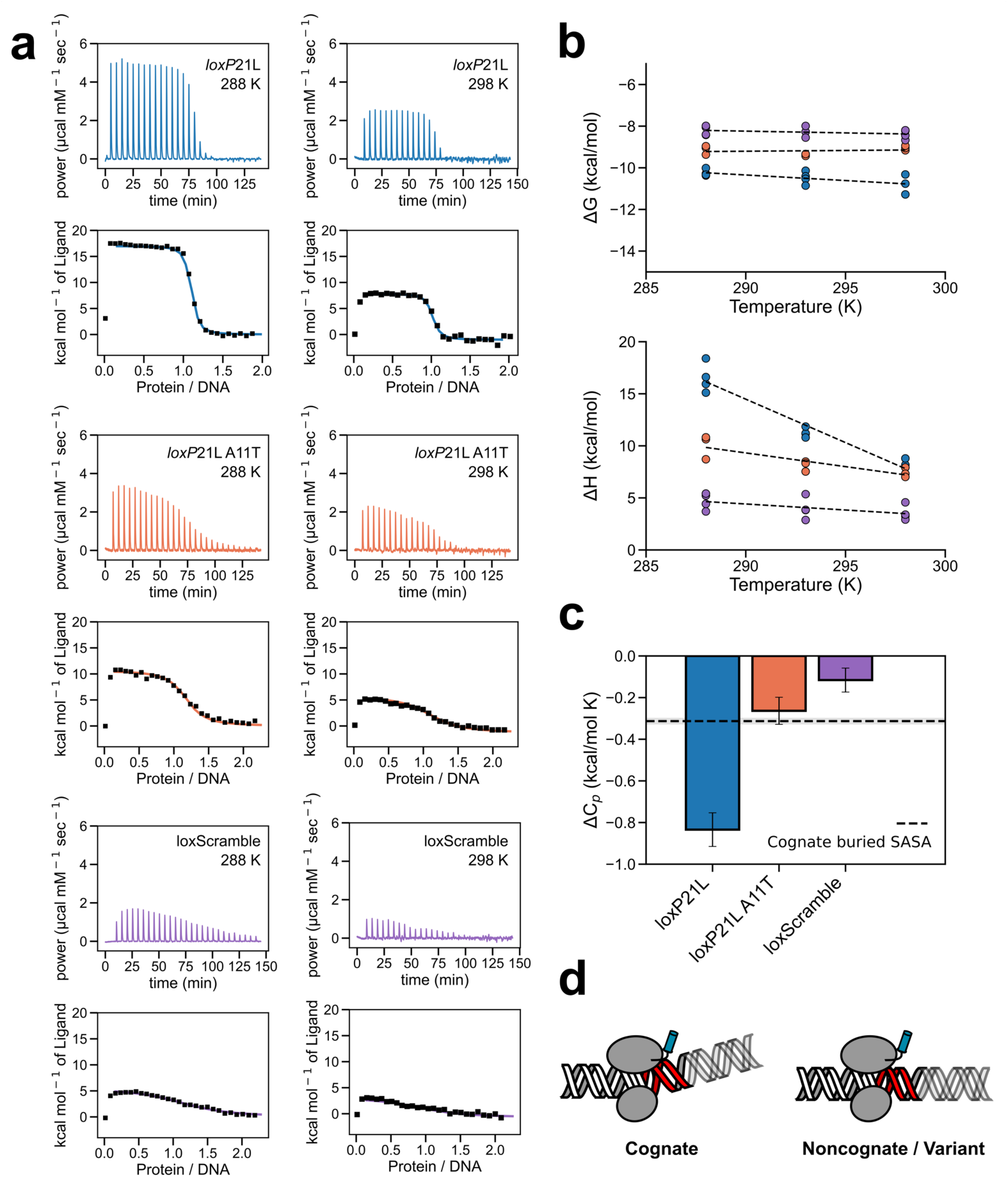
Non-contact positions in the *loxP* half-site define thermodynamic signature of specific recognition. (a) Representative thermograms for *loxP*21L (top), *loxP*21L A11T (middle), and loxScramble (bottom) reveal temperature dependent enthalpy of binding. (b) Resulting thermodynamic parameters from fitting titrations to single site binding models. Each point represents the fit value from individual replicates. (Top) Free energy of binding (ΔG) for each oligo is relatively temperature-independent, while (bottom) the oligos exhibit quite different temperature-sensitive binding enthalpies (ΔH). (c) ΔC_p_ from linear temperature dependence of ΔH. Uncertainties were estimated from the standard error of the fit. The dashed line is the ΔC_p_ predicted from buried surface area in PBD 7RHY. (d) Model for coupled DNA bending and large ΔC_p_ for specific binding to cognate DNA, and smaller value for non-cognate oligos.

The contribution to ΔC_p_ from changes in solvation was estimated from analysis of surface area burial in the cryo-EM model of monomeric Cre bound to a *loxP* half site (PDB 7RHY^9^). Contact surface area calculations reveal burial of 4371 Å^2^ of solvent exposed surface area (SASA), with 54% polar and 46% nonpolar surface area.^31^ Using an empirical model for the contribution of changes in solvation at polar and nonpolar surfaces,^30^ ΔC_p_ = (0.32 +/- 0.04 kcal/mol)·ΔA_np_ - (0.14 +/- 0.04 kcal/mol)·ΔA_p_, yields an estimate of -310 ± 10 cal mol^-1^ from solvent exclusion in the protein-DNA interface. This value is much smaller than observed for binding of Cre to *loxP*21L, within error of the value for *loxP*21L A11T, and greater than that observed for loxScramble (Figure 3 c).

For the cognate *loxP*21L interaction with Cre, the excess ∼500 cal mol^-1^ K^-1^ is consistent with substantial structural rearrangements in both protein and DNA that are coupled to site recognition. The much smaller negative ΔC_p_ values obtained for the A11T and Scramble variants suggest that these lower affinity interactions do not bury as much surface area upon binding or are not coupled to sizeable conformational changes.

### NMR reveals characteristic spectral signatures of protein conformational changes coupled to loxP recognition

To evaluate whether the unique thermodynamic signatures of Cre binding to the cognate and variant half-sites could be attributed to protein conformational changes, we compared NMR spectra of [U-^15^N]-Cre bound to *loxP*21L with those bound to *loxP*21L A11T and loxScramble half-sites (Figure 4). The spectrum of Cre bound to *loxP*21L A11T overlays well with that of Cre bound to the cognate *loxP*21L, whereas the Cre bound to loxScramble is quite different, resembling that of Cre bound to a noncognate DNA substrate (NCD1) (see also Figure S3). Spectral similarities were quantified using a nearest neighbor distance score which reciprocally measures the median difference in chemical shift for signals in a test spectrum with the signals in a corresponding reference spectrum, wherein lower scores indicate similarity. Qualitatively and quantitatively, the spectra of Cre bound to two noncognate DNA substrates with no similarity to the *loxP* RBE (NCD1 and NCD2^32^) are highly similar (median of 13 ppb), as are the spectra of Cre bound to left and right half sites (median of 18 ppb). Spectra indicate that Cre binds to *loxP*21L A11T similarly to the *loxP*21L and *loxP*21R oligos (16 and 26 ppb respectively). Conversely, when bound to the loxScramble oligonucleotide, which retains the direct-readout nucleotides, but not the non-contacted positions, Cre adopts a structure more similar to that bound to noncognate DNA substrates (18 and 23 ppb, compared to 52 and 53 ppb for cognate sequences). These spectra underscore the importance of non-contacted positions in the RBEs as important structural determinants of Cre-*loxP* half-site recognition.

**Figure 4.**
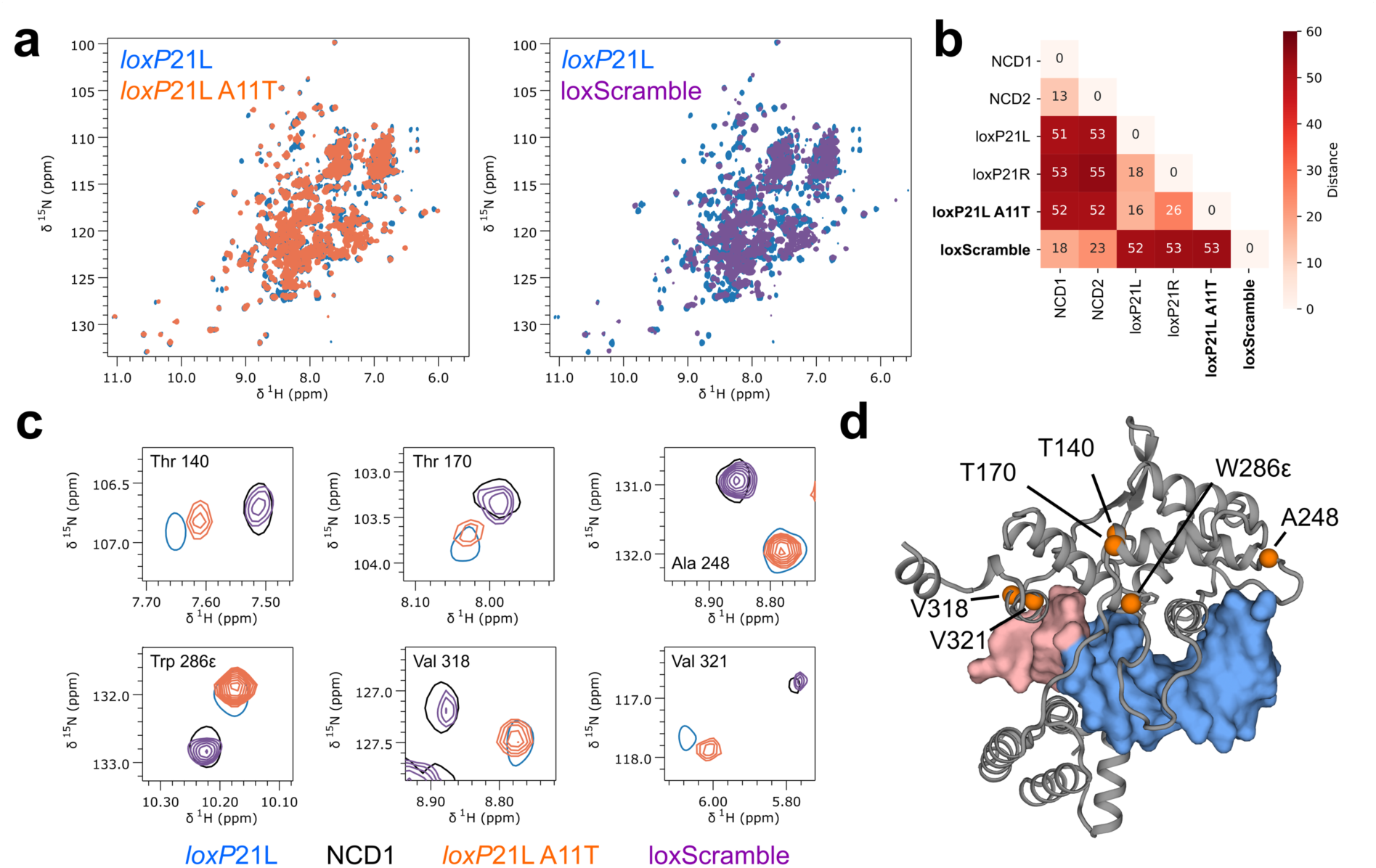
NMR spectra reveal structural role for non-contact positions in recognition. (a) Overlay of TROSY-HSQC spectra of [U-^15^N] Cre bound to *loxP*21L (blue) or its A11T (orange) or loxScramble variants (purple). (b) Pairwise spectral similarities expressed as the median difference in positions between overlaid NMR signals in parts per billion (ppb). (c) Subsets of 2D spectra of Cre bound to *loxP*21L (blue, shown as single contour), *loxP*21L A11T (orange), loxScramble (purple), and to noncognate DNA (black, shown as single contour). Assignments of resolved resonances are from the free protein. (d) Model of Cre bound to a cognate half site (chain A, 2HOI)^16^ with residues in panel c shown as orange spheres; the RBE is blue and spacer nucleotides are red. Spectra reveal similar structures for Cre bound to *loxP*21L and A11T, whereas Cre bound to the half-site mutated at all non-contact positions, loxScramble, resembles that bound to a random sequence, NCD1.

### Mutations in the loxP spacer perturb synaptic assembly

We used mass photometry to assess the effect of spacer nucleotides on dimerization and synaptic assembly. While the measured thermodynamics of monomeric Cre binding to *loxP*21L G16A exhibited no significant difference from the WT *loxP*21L sequence (Figure 2), recombination assays show the point mutation results in a 50% reduction in recombination efficiency.^13^ Similarly, prior studies showed that a *loxP* sequence containing a spacer symmetrized to the right half (*loxSR*) exhibits reduced recombination efficiency^26^ and weakened synapsis^10^, despite Cre retaining high affinity binding to half-site oligonucleotides containing that sequence (Figure 2).

The observations that these sequences matter in the context of recombination suggests the mutations affect steps after RBE recognition, such as dimerization and synapsis.

Distributions of Cre-DNA complexes detected by mass photometry revealed strong influence of spacer sequence on assembly of synaptic complexes. The WT *loxP* sequence, *loxP* G16A, and *loxSR* (Table S4) were mixed with the catalytically inactive but synapsis-competent variant, Cre K201A,^10^ at various concentrations to observe changes in oligomeric distributions. At DNA concentrations above the expected 14 nM K_D_ of synapsis^10^ (50 nM DNA, 100 nM Cre K201A, Figure 5 a), we observe three populations indicative of monomeric, dimeric, and tetrameric Cre-*loxP* complexes. These same three states are observed with *loxP* G16A, but the relative population of the tetrameric complex is decreased (Figure 5 b). For *loxSR*, only states corresponding to the monomeric and dimeric complexes are observed. Experiments at lower concentrations were used to assign the identity of the states (Figure S4). These results corroborate prior findings and suggest that mutations in the spacer that reduce recombination efficiency do so, at least in part, by destabilizing the active tetrameric complex.

**Figure 5.**
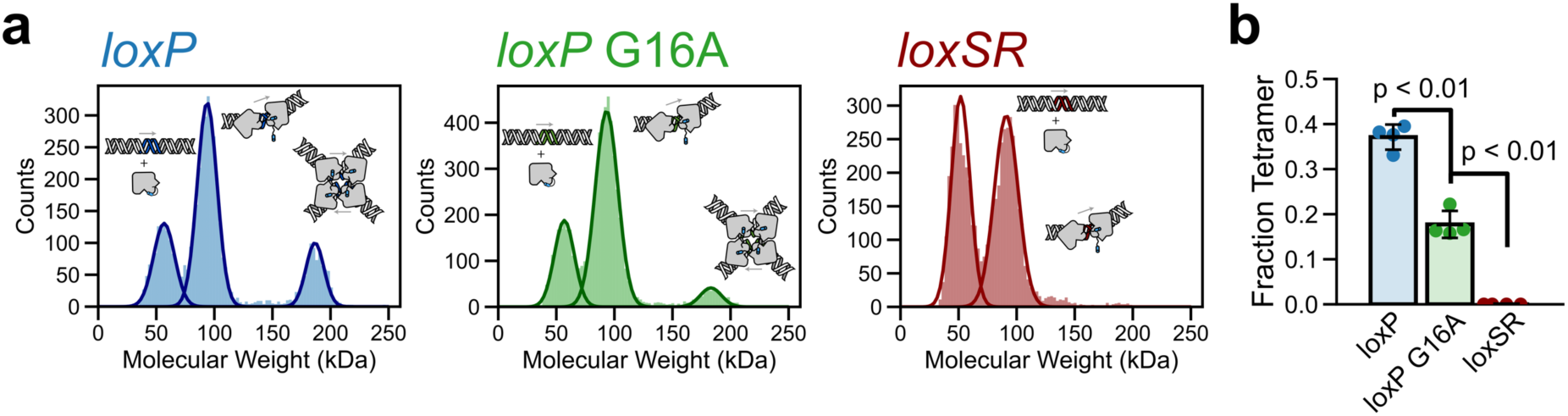
Noncontacted positions in the *loxP* spacer affect dimerization and synapsis of Cre-*loxP* complexes. Population distributions of *Cre-loxP* complexes were measured using mass photometry using catalytically inactive Cre-K201A and DNA oligonucleotides comprising two RBEs and an 8-bp spacer. Complexes of Cre-*loxP* (blue), Cre-*loxP* G16A (green), and Cre-*loxSR* (red) were formed by mixing components at 1:1 Cre to RBE ratios. Resulting histograms were fit with a Gaussian mixture model to estimate relative populations of each oligomeric state (indicated schematically) and corrected for mass-dependent differences in particle diffusion. Bar graphs represent the mean and standard deviation of the tetrameric fraction for each DNA substrate with points indicating each replicate. Significance was estimated using an unpaired t-test. Cre readily formed tetramers with *loxP*, while those were disfavored by the G16A spacer mutation, and not observed at all with the right-side symmetrized spacer *loxSR*.

## Discussion

In this work, we assessed the role of noncontacted positions in DNA on indirect readout during site recognition and assembly of higher order Cre-*loxP* complexes. We used sequence-based predictions of DNA flexibility to design variants of the *loxP* recognition sites that retained cognate nucleotides at contact positions but are expected to be resistant to protein-induced conformational changes. Titration calorimetry with these half-site substrates revealed that Cre binds these stiffer half-site substrates with reduced affinity but less unfavorable binding enthalpy. The large negative ΔC_p_ of Cre binding to cognate half-sites, but not to variant half-sites suggests that the unfavorable enthalpy of cognate binding is due to favorable deformations in the *loxP* half site substrate. NMR spectra of the protein bound to these DNA half-sites revealed that despite retaining identity at contacted positions, alteration of noncontacted positions produces structures resembling non-specific DNA binding. Additionally, we found that a single point mutation (G16A) in the *loxP* spacer, previously shown to affect recombination efficiency, destabilized higher order assemblies without affecting half-site recognition. These observations advance our understanding of the role of DNA flexibility in both recognition and assembly.

### Favorable DNA Deformations Contribute to Specificity

High resolution structures of Cre-*loxP* complexes indicate that Cre induces deformations in cognate *loxP* half site DNA, resulting in minor groove compression and DNA bending at a TATA-track.^3,9,11,33^ Site-specific high-affinity binding to *loxP* features a large unfavorable binding enthalpy (ΔH) characteristic of the energetic cost of deforming DNA substrates upon protein binding.^21,28,34,35^ To better characterize binding-coupled conformational changes in *loxP* half site variants we quantified ΔC_p_ from the temperature dependence of ΔH.^29,30^ For the cognate *loxP*21L half site, the measured ΔC_p_ is much larger than what is predicted based on a rigid body association model, consistent with coupled conformational changes.^9^ In contrast, the measured ΔC_p_ for *loxP*21L A11T does not exceed that predicted from buried surface area, suggesting that DNA deformations are not coupled to binding. The A11T mutation beaks the TATA track, removing the flexible dT-dA base step and presumably prevents DNA bending necessary for high-affinity recognition. The much smaller ΔC_p_ for Cre binding to loxScramble suggests that its resistance to deformation results in a less intimate interaction, despite retaining the nucleotides involved in direct readout.

*Protein conformational changes are coupled to recognition*.

Given its flexibly-linked two-domain structure, conformational changes in Cre are expected upon forming a C-clamp around a *loxP* RBE. The C-terminus of Cre adopts an autoinhibited conformation that occludes the DNA binding surface of the catalytic domain, yet is extended upon cognate *loxP* binding^8,32^. This C-terminal helix (αN) is also implicated in promoting assembly of Cre-*loxP* oligomers and activating Cre protomers for strand cleavage^9,10^. Protein amide NMR spectra indicate that Cre adopts a similar conformation when bound to the cognate half-site and to the A11T variant (Figure 4), despite >10-fold weaker affinity and smaller ΔC_p_ (Figures 2 and 3). In contrast, despite retaining positions involved in direct readout, the spectrum of Cre in complex with the loxScramble half-site resembles that of a non-specific complex (Figure 4).^21^ These observations are consistent with a model in which only high affinity, cognate binding is coupled to unlocking the autoinhibited conformation, with implications for assembly of higher order complexes.

### Noncontacted positions in the spacer perturb dimerization and synapsis

Experiments with longer DNA substrates containing both RBEs illustrated the importance of indirect readout in assembly of higher-order recombigenic Cre-*loxP* complexes. The specificity of Cre recombinase is not only predicated on the ability of a monomer to recognize the cognate RBEs, but also the cooperative assembly of active Cre_4_-*loxP*_2_ complexes through the formation of protein-protein interactions along and across bent DNA duplexes.^9,10,36^ While half-site substrates containing the *loxP* G16A and loxSR mutations are recognized by Cre with equivalent thermodynamics as the cognate sequences, full-length substrates with these mutations in the spacer region exhibit reduced recombination.^26^ Our mass photometry experiments showed that a single rigidifying mutation in the *loxP* spacer (G16A) significantly reduces the population of Cre_4_-*loxP*_2_ complexes, while a symmetric right spacer abolished synapsis of Cre_2_-*loxP* complexes under the tested conditions, consistent with their reported effects on recombination.^13^ These results demonstrate underscore the importance of non-contacted positions in the *loxP* spacer in defining stability of synaptic complexes and subsequent recombination.

## Conclusions

These experiments highlight the role of DNA flexibility in indirect readout and specificity of Cre-*loxP* recognition. Detailed calorimetric analysis revealed that direct base-specific contacts to the *loxP* sequence do not make dominant contributions to high-affinity binding, rather Cre recognizes target *loxP* half sites through coupled conformational changes in the form of DNA deformations. We show that noncontacted positions in the *loxP* sequence drive site-specific recognition, and that variants with altered recombinase binding elements exhibit thermodynamic and spectral signatures comparable to noncognate DNA. We also observe that mutations in the *loxP* spacer can reduce recombination efficiency by destabilizing the synaptic complex rather than perturbing monomer recognition. This work advances our understanding of site selection by Cre recombinase and reveal thermodynamic and spectroscopic signatures of site recognition that could be exploited in the engineering of Cre variants with altered site specificities.

## Methods

### Protein expression and purification

A pET21A plasmid encoding for the wild-type Cre enzyme (P06956) was provided by Gregory Van Duyne (University of Pennsylvania Philadelphia, PA). The plasmid encoding Cre K201A was described previously^9,32^. Electrocompetent *E. coli* T7 express LysY/Iq cells (New England Biolabs) were transformed using a 1 mm cuvette and plated on LB agarose plates supplemented with 0.1 mg/mL carbenicillin for antibiotic selection and grown overnight at 37 °C. A single colony was picked from the antibiotic selection plate and was used to inoculate to a liquid 50 mL LB media culture supplemented with 0.1 mg/mL carbenicillin. This culture was grown overnight at 37 °C shaken at 220 rpm.

Following overnight growth, 10 mL of liquid culture was transferred to 1 L of either rich LB media or M9 minimal media^32^ supplemented with 1 g/L ^15^NH_4_ as the sole nitrogen source for protein overexpression. Liquid cultures were additionally supplemented with 0.1 mg/mL carbenicillin for antibiotic selection and incubated at 37 °C shaking at 220 rpm. Once the optical density at 600 nm reached 0.6-0.8, 400 μL of 1 M IPTG (Isopropyl β-D-1-thiogalactopyranoside) was added to induce protein expression and the cultures were allowed to incubate for an additional 4 hours. Following the induction period, cells were pelleted by centrifugation at 4225 x g, liquid media was decanted, and pellets were frozen at -80 °C.

Frozen cell pellets containing expressed protein were thawed on ice and resuspended in 35 mL of buffer A (40 mM Tris pH 7.0, 100 mM NaCl) supplemented with 1 mM EDTA and one tablet of miniComplete EDTA free protease inhibitor (Roche).

Cells were lysed by sonication on ice (Qsonica, two cycles of 50% amplitude for 5 minutes, 5 seconds on and 5 seconds off) and the insoluble fraction was pelleted by centrifugation at 26891 x g for 45 minutes at 4 °C. Supernatant was syringe filtered using a 0.2 μm filter (Pall) and first purified using cation exchange chromatography using a 5 mL SPFF column (Cytiva) eluted with a linear salt gradient (60 mL, 40 mM Tris pH 7.0, 100 mM to 1 M NaCl). Fractions containing Cre protein were isolated and diluted 1:4 in buffer A to reduce the ionic strength and were further purified by affinity chromatography with a 5 mL heparin column (Cytiva) eluted with a linear salt gradient (90 mL, 40 mM Tris pH 7.0, 100 mM to 1.5 M NaCl). Eluted protein was concentrated to 2 mL before loading onto size exclusion using a 120 mL S75pg column (Cytiva) in S75 buffer (40mM Tris pH 7.0, 500 mM NaCl, 5 mM DTT). Purity was ensured to be >95% using an SDS-PAGE gel.

### DNA design and purification

Variant *loxP* half-site sequences were first designed by predicting the effect of mutations at noncontacted positions on the DNA minor groove width. Sequences were chosen that perturb the predicted minor groove expansion and compression between positions 12-18 (Figure 1a, Figure S1), as expansion and compression at these positions is also observed in the structure of Cre bound to a *loxP* half-site. Resulting sequences were then compared to TRX score to assay the effect of the mutation on predicted DNA flexibility.

Single-strand DNA oligos (Integrated DNA Technologies) were resuspended in miliQ H_2_O to 100 μM and concentrations were confirmed by nanodrop (Thermo Scientific). Resuspended top and bottom strand oligos were combined in equimolar amounts in annealing buffer (10 mM Tris pH 7.0 and 100mM NaCl) and heated at 95 °C for 10 minutes. The water bath was then turned off and the oligos slowly cooled overnight to allow for proper annealing. Annealed DNA duplexes were purified to remove ssDNA contamination by anion exchange chromatography using a QHP column (Cytiva) with a linear buffer gradient (10 mM Tris pH 7.0, 25 mM to 1 M NaCl). Purified dsDNA was ensured by analyzing chromatography fractions on a 2% (w/v) agarose gel impregnated with Sybr Safe dye (Invitrogen) for visualization.

### Isothermal titration calorimetry

WT Cre and DNA substrates were dialyzed overnight at 4 °C into 1 L of ITC buffer (20mM HEPES pH 7.0, 250 mM NaCl) using either 6-8 kDa MWCO dialysis tubing (Spectrum Laboratories) or 3.5 kDa MWCO dialysis cassettes (Thermo Scientific) in the same container to limit heats of dilution. Protein and DNA concentrations were measured after dialysis by measuring absorbance at 280 and 260 nm respectively (ε_280,WT_ = 48,930 M^-1^ cm^-1^, ε_260,loxP21L_ = 333,780 M^-1^ cm^-1^, ε_260,loxP21R_ = 334,427 M^-1^ cm^-1^, ε_260,loxP21L A11T_ = 330,581M^-1^ cm^-1^, ε_260,loxP21L G16A_ = 341,315 M^-1^ cm^-1^, and ε_260,loxScramble_ = 341,074 M^-1^ cm^-1^). Protein was loaded into the syringe at concentrations ranging from 30-150 μM and DNA was loaded into the cell at concentrations ranging from 3-15 μM. ITC experiments were run using a VP-ITC (Malvern Panalytical) at temperatures ranging from 15-25°C, reference power of 5 μcal/sec. Injection volumes were set to 10 μL with an inter-injection delay of 300 sec, and a stirring speed of 307 rpm. Baselining and integration of the thermograms was completed using NITPIC^37^ and titrations were fit using ITCSIMLIB^38^ to a single-site binding model and fit parameters and uncertainties for each titration were determined by bootstrapping with 1000 iterations. Changes in polar and nonpolar solvent accessible surface area were computed for the structure of monomer Cre bound to a *loxP* half site substrate (PDB 7RHY) using dr-SASA (github.com/nioroso-x3/dr_sasa_n)^31^.

### Nuclear magnetic resonance

Protein-DNA complexes were formed by mixing [U-^15^N]-Cre with two-fold excess DNA substrate in high salt NMR buffer (20 mM HEPES pH 7.0, 500 mM NaCl, 5 mM DTT). The mixture was then dialyzed stepwise at 4 °C, each step a 100-fold dilution for 8 hours into NMR buffer with reduced ionic strength (500◊ 200◊ 100◊ 25 mM NaCl) in 6-8 kDa MWCO dialysis tubing (Spectrum Laboratories). Complexes were then concentrated by centrifugation using a 3 kDa spin cutoff filter to an approximate final concentration of 300 mM Cre. 250 μL of complex were supplemented with 10% (v/v) D_2_O and 225 μM DSS (2,2-dimethyl-2-silapentane-5-sulfonate) for spin locking and chemical shift referencing respectively in 3 mm NMR tubes. ^1^H-^15^N TROSY-HSQC spectra were recorded using a Bruker Avance 800 MHz spectrometer equipped with a triple-resonance inverse (TXI) cryoprobe. Spectra were collected with 40-48 scans, 2048 x 256 complex points, and a spectral width of 21.85 x 36.00 ppm centered at 4.70 and 116 ppm for the ^1^H and ^15^N dimensions respectively. Data were processed using NMRFx Analyst^39^ (nmrfx.org). Spectral similarity scores were computed from:

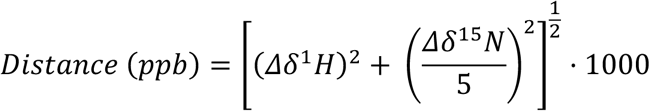

### Mass photometry

Purified Cre K201A, *loxP*38, *loxP*38 G16A, and *loxSR* were dialyzed overnight against buffer containing 20 mM HEPES pH 7.0 and 100 mM NaCl for a 3:1000 dilution.

Protein-DNA complexes were allowed to equilibrate for more than two hours before experimentation. Borosilicate slides were cleaned in triplicate with alternating washes of miliQ-H_2_O and isopropanol, finishing with H_2_O before drying with nitrogen gas. Mass photometry measurements were completed using a Refeyn TwoMP instrument with buffer-free focusing and direct loading of samples for measurement. Calibration of radiometric contrast to molecular weights was completed using standards with known molecular weights of 86, 172, 258 and 344 kDa from the Refyn MassFerence P1 calibrant diluted in the buffer above. Movies measuring radiometric contrast of scattering events were recorded for 60 seconds and contrast values were converted to molecular weights using the calibration curve. Data were exported as individual molecular weight values and histograms were fit using a Gaussian mixture model using the SciPy and Scikit-learn python libraries. Estimated populations were calculate from the relative weight of the gaussian fit, normalized to molecular weight dependent differences in diffusion for each species (D ∝ MW^-1/3^). Code available at GITHUB. Fraction of Cre monomers that assemble into tetrameric complexes was estimated as:

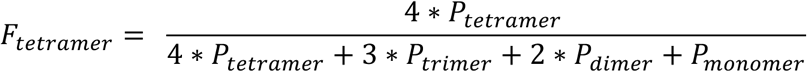

## Acknowledgements

This work was supported by NIH R01 GM122432 (to M.P.F). J.S.M was supported by NIH T32 GM144293. MJB was supported by T32 GM141955. R.M.B. received support from The Ohio State University’s Summer Research Opportunities Program (SROP). Access to calorimetry instrumentation was provided by the Department of Chemistry and Biochemistry; NMR and mass photometry data were recorded at the Campus Chemical Instrument Center. Analysis and management of NMR data was enabled by NMRbox: National Center for Biomolecular NMR Data Processing and Analysis, a Biomedical Technology Research Resource (BTRR), which is supported by NIH grant P41GM111135 (NIGMS).

