## Supporting Information for "Thermodynamics of Indirect Readout in Cre-*loxP* Recognition"

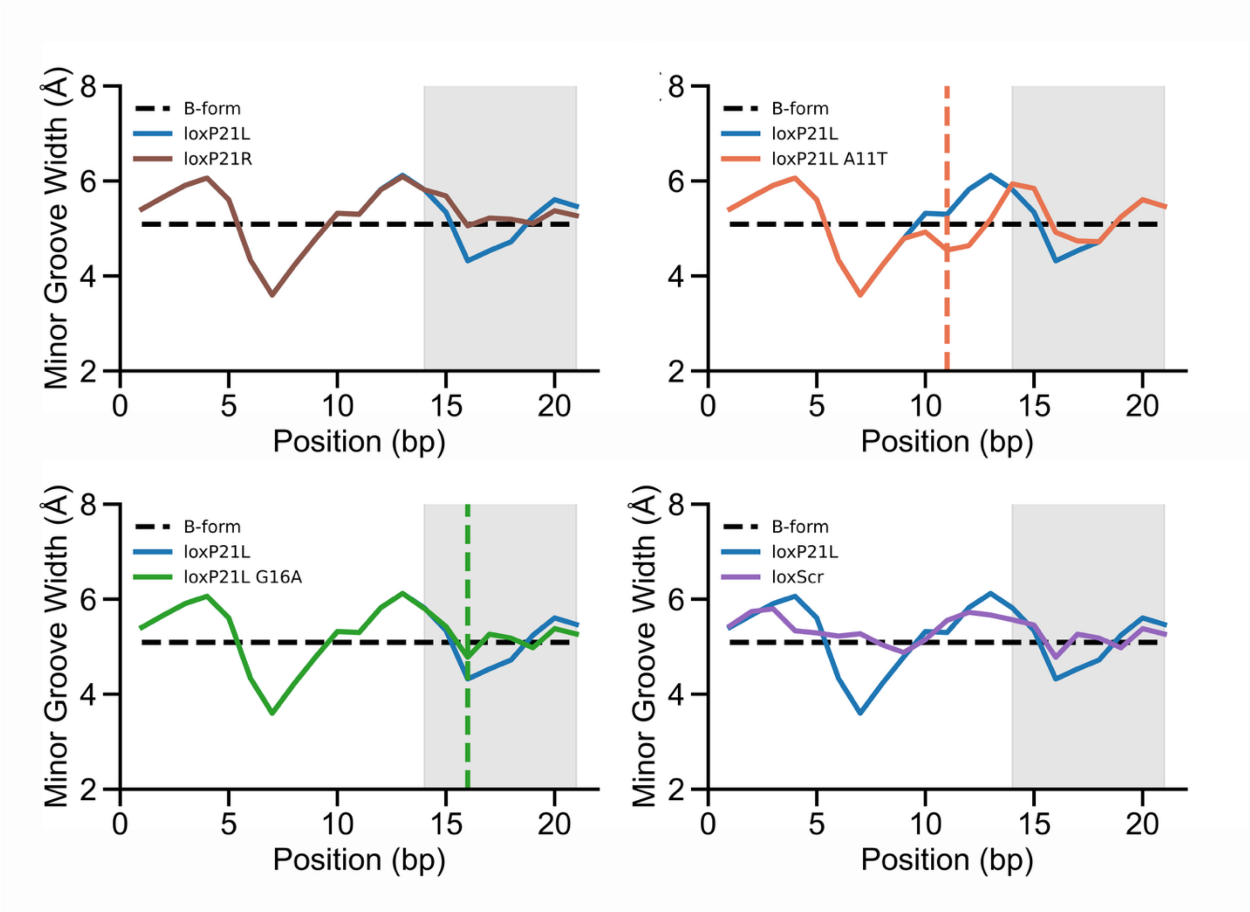

**Supplemental Figure 1.** Predicted minor groove width for *loxP* half site variants using the DEEP DNAshape webserver (https://deepdnashape.usc.edu/). Shaded regions indicate the position of the *loxP* spacer sequence and vertical dashed lines indicate the position of single point mutations. Black dashed lines indicate minor groove width for B-form DNA.

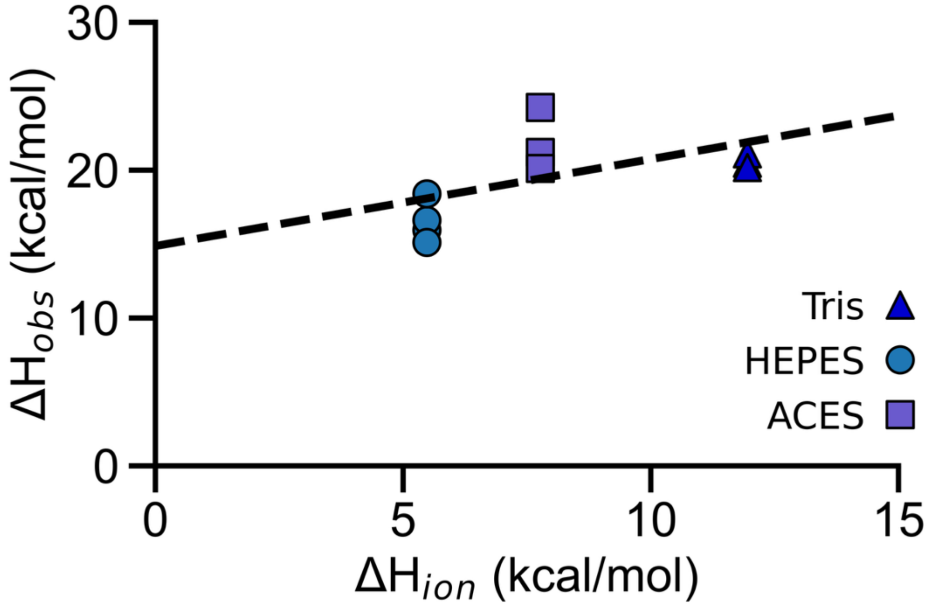

**Supplemental Figure 2. Effect of buffer ionization on enthalpy of Cre-loxP half site binding.** The observed enthalpy ($\Delta H_{obs}$) of WT Cre binding *loxP*21L substrates was measured at 15 °C in three buffers exhibiting different enthalpies of ionization ($\Delta H_{ion}$). Fitting this relationship yields a buffer corrected enthalpy of binding of +14.86 kcal/mol and 0.6 ± 0.3 ionization events at pH 7.0.

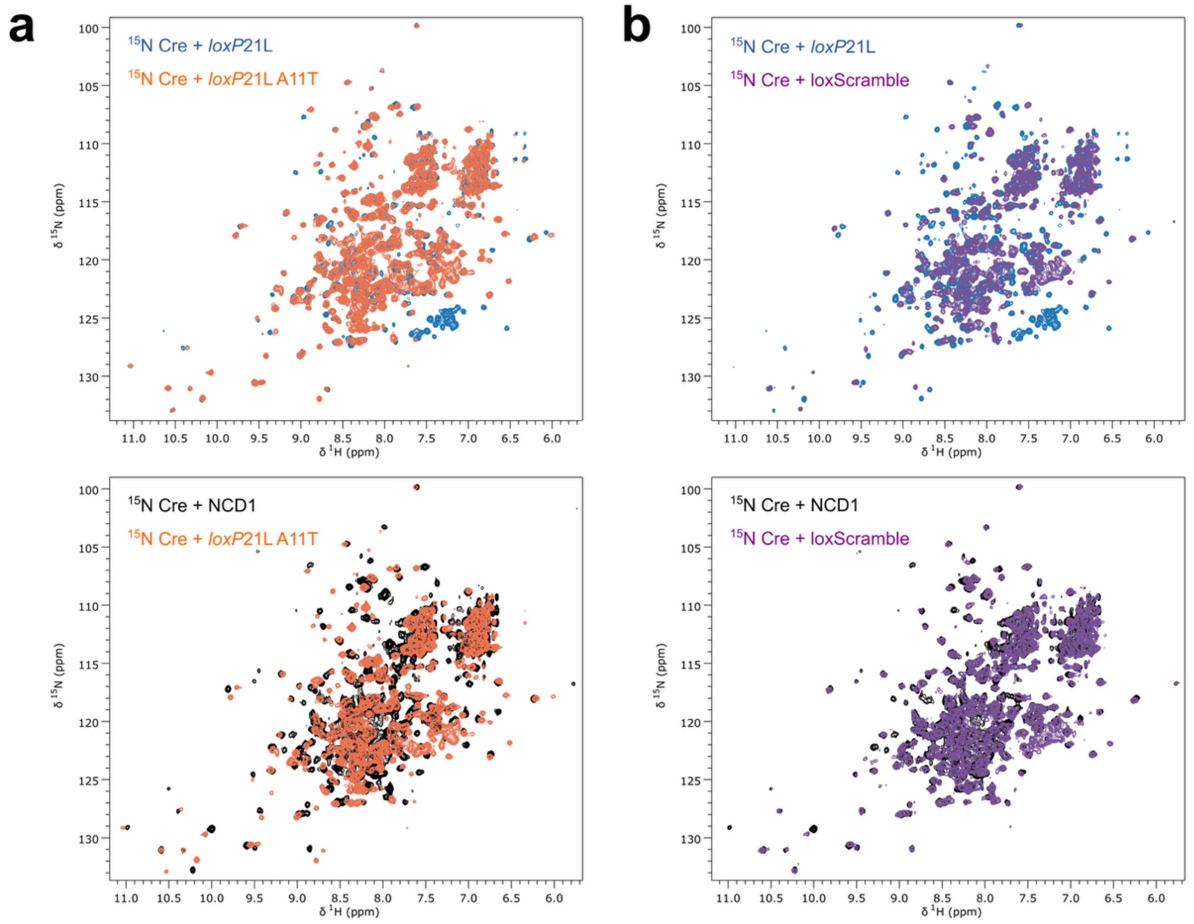

**Supplemental Figure 3. TROYS-HSQC spectra of [U-$^{15}$N] Cre in complex with DNA.** Spectra of Cre mixed with excess cognate *loxP*21L (blue) or noncognate DNA (black) were compared to spectra of Cre mixed with excess loxP21L A11T (orange) or loxScramble (purple). Qualitatively, it appears that the spectrum of Cre-A11T overlays more closely with loxP21L whereas the spectrum of Cre-loxScramble overlays more closely with noncognate DNA.

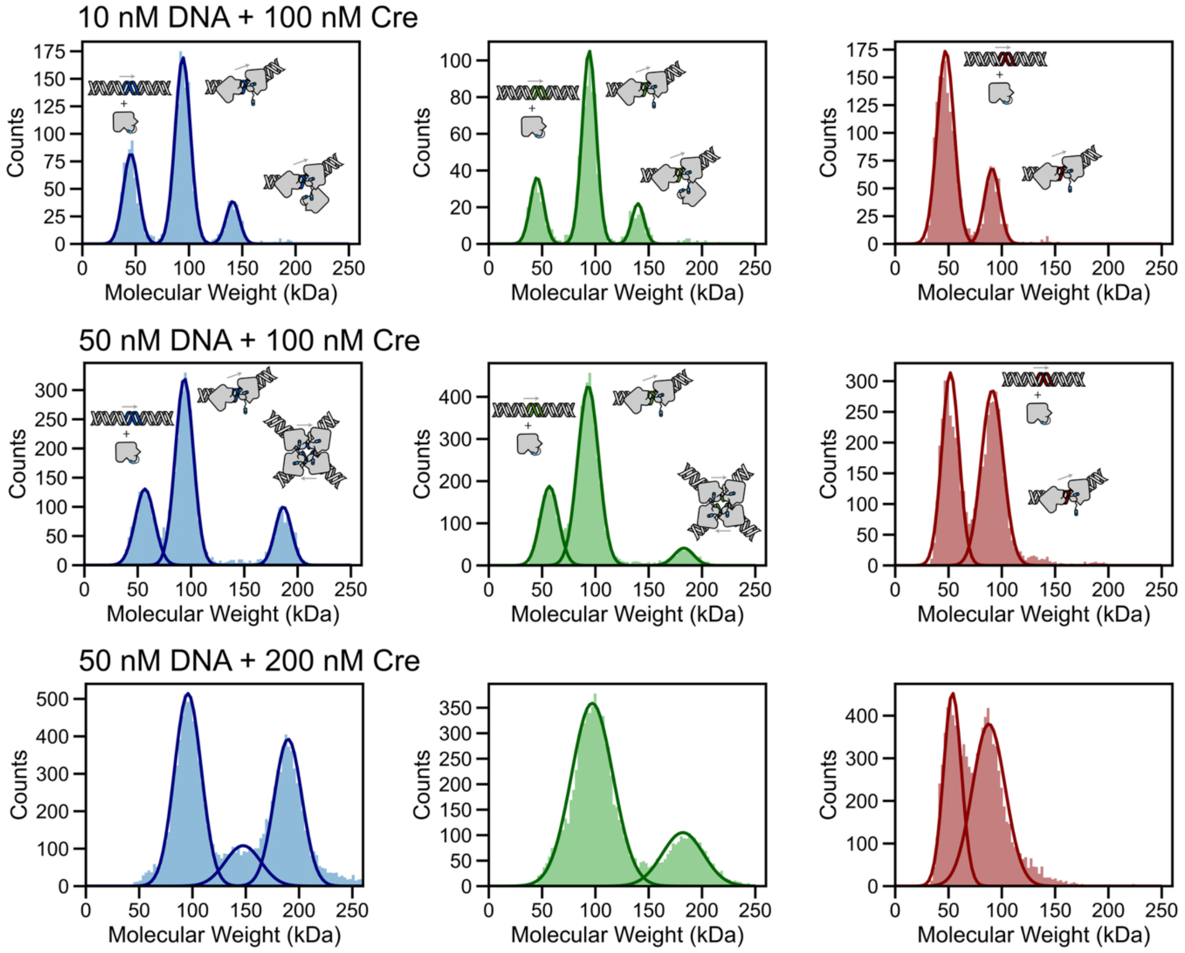

**Supplemental Figure 4. Mutations in the loxP spacer perturb synapsis.** Population distributions for assembly intermediates of Cre- *loxP* (blue), *loxP* G16A (green), and loxSR (red) complexes at varying protein and DNA concentrations. We observe a ~45 kDa species that corresponds to free Cre or the monomer bound to loxP, a 94 kDA species that corresponds to a Cre2-loxP dimer, a 140 kDa species that corresponds to a Cre3-loxP complex, and a 190 kDa species that corresponds to the tetrameric complex. Populations were fit using a gaussian mixture model.

| DNA Substrate | Run | [Cell] (μM) | n | +/- | ΔG (kcal/mol) | +/- | ΔH (kcal/mol) | +/- | Kd (nM) | +/- |
|---|---|---|---|---|---|---|---|---|---|---|
| loxP21L | 1 | 15 | 1.05 | 0.004 | -10.28 | 0.15 | 15.94 | 0.14 | 15.80 | 0.23 |
| loxP21L | 2 | 15 | 1.07 | 0.005 | -10.01 | 0.12 | 16.61 | 0.15 | 25.33 | 0.30 |
| loxP21L | 3 | 10 | 0.97 | 0.010 | -10.37 | 0.44 | 18.40 | 0.44 | 13.50 | 0.38 |
| loxP21L | 4 | 4 | 1.16 | 0.010 | -10.33 | 0.33 | 15.11 | 0.33 | 14.48 | 0.32 |
| | | Mean | 1.06 | 0.01 | -10.25 | 0.26 | 16.52 | 0.27 | 17.28 | 0.31 |
| | | Stdev | 0.08 | | 0.16 | | 1.40 | | 5.45 | |

| DNA Substrate | Run | [Cell] (μM) | n | +/- | ΔG (kcal/mol) | +/- | ΔH (kcal/mol) | +/- | Kd (nM) | +/- |
|---|---|---|---|---|---|---|---|---|---|---|
| loxP21L T7A | 1 | 4 | 1.26 | 0.02 | -10.11 | 0.18 | 13.98 | 0.28 | 27.06 | 0.54 |
| loxP21L T7A | 2 | 4.5 | 1.24 | 0.01 | -9.57 | 0.11 | 14.24 | 0.21 | 68.06 | 1.01 |
| loxP21L T7A | 3 | 4 | 1.34 | 0.01 | -9.69 | 0.10 | 14.08 | 0.18 | 55.82 | 0.72 |
| | | Mean | 1.28 | 0.01 | -9.79 | 0.13 | 14.10 | 0.22 | 50.31 | 0.76 |
| | | Stdev | 0.05 | | 0.28 | | 0.13 | | 21.04 | |

| DNA Substrate | Run | [Cell] (μM) | n | +/- | ΔG (kcal/mol) | +/- | ΔH (kcal/mol) | +/- | Kd (nM) | +/- |
|---|---|---|---|---|---|---|---|---|---|---|
| loxP21L A11T | 1 | 5 | 1.16 | 0.02 | -9.04 | 0.09 | 10.63 | 0.16 | 137.47 | 1.31 |
| loxP21L A11T | 2 | 5 | 1.10 | 0.04 | -9.37 | 1.20 | 8.71 | 0.39 | 77.07 | 9.87 |
| loxP21L A11T | 3 | 5 | 1.09 | 0.01 | -8.96 | 0.08 | 10.82 | 0.16 | 156.74 | 1.33 |
| | | Mean | 1.11 | 0.02 | -9.12 | 0.45 | 10.05 | 0.24 | 123.76 | 4.17 |
| | | Stdev | 0.04 | | 0.22 | | 1.17 | | 41.56 | |

| DNA Substrate | Run | [Cell] (μM) | n | +/- | ΔG (kcal/mol) | +/- | ΔH (kcal/mol) | +/- | Kd (nM) | +/- |
|---|---|---|---|---|---|---|---|---|---|---|
| loxP21L G16A | 1 | 3 | 1.23 | 0.01 | -10.50 | 0.16 | 14.54 | 0.22 | 10.73 | 0.16 |
| loxP21L G16A | 2 | 3 | 1.20 | 0.01 | -10.59 | 0.20 | 15.10 | 0.26 | 9.18 | 0.16 |
| loxP21L G16A | 3 | 3 | 1.16 | 0.01 | -10.63 | 0.16 | 15.19 | 0.21 | 8.46 | 0.11 |
| | | Mean | 1.20 | 0.01 | -10.57 | 0.17 | 14.94 | 0.23 | 9.45 | 0.14 |
| | | Stdev | 0.03 | | 0.07 | | 0.35 | | 1.16 | |

| DNA Substrate | Run | [Cell] (μM) | n | +/- | ΔG (kcal/mol) | +/- | ΔH (kcal/mol) | +/- | Kd (nM) | +/- |
|---|---|---|---|---|---|---|---|---|---|---|
| loxScr | 1 | 10 | 1.12 | 0.04 | -8.06 | 0.12 | 4.43 | 0.14 | 765.33 | 11.78 |
| loxScr | 2 | 10 | 1.18 | 0.02 | -7.99 | 0.07 | 5.22 | 0.09 | 861.95 | 7.12 |
| loxScr | 3 | 5 | 1.03 | 0.04 | -8.42 | 0.12 | 5.42 | 0.17 | 403.60 | 5.51 |
| loxScr | 4 | 4.5 | 1.06 | 0.04 | -8.19 | 0.13 | 3.67 | 0.13 | 604.43 | 9.45 |
| | | Mean | 1.10 | 0.03 | -8.16 | 0.11 | 4.68 | 0.14 | 658.83 | 8.47 |
| | | Stdev | 0.07 | | 0.19 | | 0.80 | | 200.59 | |

**Supplemental Table 1. Thermodynamic parameters obtained from fitting titrations of mutant loxP half site sequences at 15°C.** Uncertainties in individual fits were estimated by bootstrapping with 1000 iterations.

| Buffer | Run | [Cell] (μM) | n | +/- | ΔG (kcal/mol) | +/- | ΔH (kcal/mol) | +/- | Kd (nM) | +/- |
|---|---|---|---|---|---|---|---|---|---|---|
| HEPES | 1 | 15 | 1.05 | 0.004 | -10.28 | 0.15 | 15.94 | 0.14 | 15.80 | 0.23 |
| HEPES | 2 | 15 | 1.07 | 0.005 | -10.01 | 0.12 | 16.61 | 0.15 | 25.33 | 0.30 |
| HEPES | 3 | 10 | 0.97 | 0.010 | -10.37 | 0.44 | 18.40 | 0.44 | 13.50 | 0.38 |
| HEPES | 4 | 4 | 1.16 | 0.010 | -10.33 | 0.33 | 15.11 | 0.33 | 14.48 | 0.32 |
| | | Mean | 1.06 | 0.01 | -10.25 | 0.26 | 16.52 | 0.27 | 17.28 | 0.31 |
| | | Stdev | 0.08 | | 0.16 | | 1.40 | | 5.45 | |

| Buffer | Run | [Cell] (μM) | n | +/- | ΔG (kcal/mol) | +/- | ΔH (kcal/mol) | +/- | Kd (nM) | +/- |
|---|---|---|---|---|---|---|---|---|---|---|
| Tris | 1 | 4 | 1.00 | 0.02 | -11.16 | 2.79 | 20.47 | 0.56 | 3.36 | 0.84 |
| Tris | 2 | 4 | 0.95 | 0.01 | -10.85 | 0.14 | 21.16 | 0.26 | 5.75 | 0.07 |
| Tris | 3 | 4 | 0.93 | 0.06 | -11.06 | 3.87 | 20.18 | 2.58 | 4.00 | 1.40 |
| | | Mean | 0.96 | 0.03 | -11.02 | 2.27 | 20.60 | 1.13 | 4.37 | 0.77 |
| | | Stdev | 0.04 | | 0.16 | | 0.51 | | 1.23 | |

| Buffer | Run | [Cell] (μM) | n | +/- | ΔG (kcal/mol) | +/- | ΔH (kcal/mol) | +/- | Kd (nM) | +/- |
|---|---|---|---|---|---|---|---|---|---|---|
| ACES | 1 | 2.5 | 1.02 | 0.02 | -11.71 | 4.00 | 21.20 | 0.81 | 1.28 | 0.43 |
| ACES | 2 | 2.5 | 1.01 | 0.03 | -11.07 | 4.10 | 24.24 | 1.05 | 3.93 | 1.45 |
| ACES | 3 | 2.5 | 1.02 | 0.01 | -10.55 | 0.10 | 20.11 | 0.19 | 9.79 | 0.10 |
| | | Mean | 1.02 | 0.02 | -11.11 | 2.73 | 21.85 | 0.68 | 5.00 | 0.66 |
| | | Stdev | 0.01 | | 0.58 | | 2.14 | | 4.35 | |

**Supplemental Table 2. Thermodynamic parameters from fits of Cre-*loxP*21L titrations with varying buffer enthalpy of ionization.** Buffers contain 20 mM of buffering agent at pH 7.0 and 250 mM NaCl. Uncertainties in individual fits were estimated by bootstrapping with 1000 iterations.

| DNA Substrate | Temperature (K) | n | +/- | ΔG (kcal/mol) | +/- | ΔH (kcal/mol) | +/- | Kd (nM) | +/- |
|---|---|---|---|---|---|---|---|---|---|
| loxP21L | 288 | 1.06 | 0.08 | -10.25 | 0.16 | 16.52 | 1.40 | 17.28 | 5.45 |
| loxP21L | 293 | 1.05 | 0.01 | -10.48 | 0.30 | 11.25 | 0.44 | 15.60 | 7.60 |
| loxP21L | 298 | 1.19 | 0.02 | -10.79 | 1.84 | 8.29 | 0.45 | 15.03 | 11.00 |
| loxP21L A11T | 288 | 1.11 | 0.04 | -9.12 | 0.21 | 10.05 | 1.16 | 123.76 | 4.17 |
| loxP21L A11T | 293 | 1.07 | 0.02 | -9.38 | 0.10 | 8.12 | 0.14 | 100.76 | 1.06 |
| loxP21L A11T | 298 | 1.11 | 0.01 | -9.05 | 0.11 | 7.42 | 0.46 | 178.08 | 33.20 |
| loxScramble | 288 | 1.10 | 0.07 | -8.16 | 0.19 | 4.68 | 0.80 | 658.83 | 200.59 |
| loxScramble | 293 | 1.02 | 0.11 | -8.27 | 0.23 | 3.98 | 1.03 | 717.08 | 279.60 |
| loxScramble | 298 | 1.02 | 0.13 | -8.38 | 0.22 | 3.53 | 0.72 | 743.98 | 251.29 |

**Supplemental Table 3. Thermodynamic parameters from fitting titrations at varying temperatures.**

Uncertainties represent the standard deviation of all replicates.

| Name | Sequence | $\varepsilon_{260}$ ($M^{-1}$ $cm^{-1}$) | Use |
|---|---|---|---|
| **loxP21L** | 5' – GCA TAA CTT CGT ATA ATG TCG – 3'<br>3' – CGT ATT GAA GCA TAT TAC AGC – 5' | 333,780 | ITC/NMR |
| **loxP21R** | 5' – GCA TAA CTT CGT ATA GCA TCG – 3'<br>3' – CGT ATT GAA GCA TAT CGT AGC – 5' | 334,427 | ITC/NMR |
| **loxP21L A11T** | 5′ – GCA TAA CTT CGT TTA ATG TCG – 3′<br>3′ – CGT ATT GAA GCA AAT TAC AGC – 5′ | 330,581 | ITC/NMR |
| **loxP21L G16A** | 5' – GCA TAA CTT CGT ATA ATA TCG – 3'<br>3' – CGT ATT GAA GCA TAT TAT AGC – 5' | 341,315 | ITC |
| **loxScramble** | 5' – GCA GAC AAG CAT ATA ATA TCG – 3'<br>3' – CGT CTG TTC GTA TAT TAT AGC – 5' | 341,074 | ITC/NMR |
| **loxP 38** | 5′ – GCA TAA CTT CGT ATA ATG TAT GCT ATA CGA AGT TAT CG – 3′<br>3′ – CGT ATT GAA GCA TAT TAC ATA CGA TAT GCT TCA ATA GC – 5′ | 602,426 | MP |
| **loxP 38 G16A** | 5′ – GCA TAA CTT CGT ATA ATA TAT GCT ATA CGA AGT TAT CG – 3′<br>3′ – CGT ATT GAA GCA TAT TAT ATA CGA TAT GCT TCA ATA GC – 5′ | 602,409 | MP |
| **loxSR** | 5′ – GGA TAA CTT CGT ATA GCA TAT GCT ATA CGA AGT TAT CC – 3′<br>3′ – CCT ATT GAA GCA TAT CGT ATA CGA TAT GCT TCA ATA GG – 5′ | 605,082 | MP |

**Supplemental Table 4: DNA extinction coefficients.** DNA sequence is colored to indicate the RBE (blue) and spacer (red). Extinction coefficients shown were used to determine DNA concentration.
